# Beta and Gamma Dynamics in Attentional Networks Predict Conscious Reports

**DOI:** 10.1101/2025.03.24.644800

**Authors:** Alfredo Spagna, Jianghao Liu, Paolo Bartolomeo

**Author notes:** **Conflict of Interest:** The authors declare that they have no competing financial interests.

## Abstract

What neural events precede conscious reports? Hemisphere-asymmetric attentional networks are causally related to conscious perception (Bartolomeo et al., 2025; Kaufmann et al., 2024), but their spectrotemporal dynamics remain unclear. Here, we used magnetoencephalography to examine brain oscillations occurring in human participants (male and female) before a near-threshold target during the cue-target period. In 67% of trials, a supra-threshold visual cue appeared near the target placeholder box, indicating that the target would appear at that location (valid condition). In the remaining 33% of trials, the target appeared at the opposite location (invalid condition). We analyzed brain oscillations, coherence, and theta-gamma phase-amplitude (PAC) coupling in 18 regions of interest involved in attentional and perception (Martín-Signes et al., 2024). Results revealed that: (1) Report of validly cued targets was preceded by early (∼58 ms post-cue) beta-band activity in the right-hemisphere superior parietal lobule. (2) Report of invalidly cued targets was preceded by late (∼166 ms post-cue) beta-band activity in the right temporo-occipital (TO) region, PAC in the right lateral visual cortex, and low gamma coherence between this region and the left temporo-parietal junction, (3) Unreported invalidly cued targets were preceded by PAC in the right TO, and by high gamma coherence between this region and the right middle frontal gyrus suggesting a pre-target bias. We show that conscious report is preceded by temporally dissociable, frequency-specific reconfigurations of right-lateralized attentional networks, with an early parietal beta-mediated orienting window and a later ventral beta- and gamma-mediated window that predict conscious reports before target onset.

**Significance statement:** Conscious perception depends not only on sensory signals, but also on how attentional networks prepare the brain in advance. Using MEG and a spatial cueing task with near-threshold targets, we show distinct right-hemisphere beta- and gamma-band dynamics predicting whether an upcoming stimulus will be reported or missed. Validly cued reports rely on early beta activity in the right superior parietal lobule, whereas invalid reports and omission errors are linked to later beta activity in the right temporo-occipital region, early theta–gamma coupling between this region and the right middle frontal gyrus, and theta-gamma phase-amplitude coupling. These findings reveal hemisphere-asymmetric spectrotemporal signatures by which attention biases predictive processing and shapes the conscious report of future stimuli, informing theories of conscious perception.

## Introduction

Spatial cues guide our behavior by helping identify crucial information and potential dangers. For example, blinking traffic lights signal the need for increased attention and rapid decision-making at intersections. Frontoparietal oscillations have been proposed as a key neural mechanism supporting conscious target processing after appearance (Nobre & Van Ede, 2023; Tosoni et al., 2023). However, the oscillatory dynamics *before* target onset remain less explored, despite this being the critical period for environmental sampling (Fiebelkorn & Kastner, 2019; VanRullen, 2016; Weisz et al., 2014; Zhou et al., 2021) and attentional allocation (Fries, 2023; VanRullen, 2018). We address this gap by investigating multiple spectrotemporal dynamics in response to predictive peripheral cues preceding near-threshold targets..

Prestimulus oscillatory activity influences whether near-threshold stimuli reach awareness by shifting the decision criterion and altering sensitivity. In particular, alpha-band dynamics have been interpreted within local functional inhibition and network-level gating frameworks, whereby oscillatory power indexes cortical excitability and shapes large-scale information routing prior to stimulus onset (Baumgarten et al., 2023; Weisz et al., 2014; Zhou et al., 2021). Spatial attention can modulate preparatory states via exogenous or endogenous mechanisms, either modifying sensory signals (Carrasco, 2011; Carrasco & Barbot, 2019; Chica, et al., 2013; Hanning et al., 2019) or biasing decision criteria (Bang & Rahnev, 2017). Exogenous cues capture attention automatically, while predictive cues engage endogenous, voluntary shifts (Asanowicz et al., 2023). Peripheral predictive cues engage both processes (Chica et al., 2011; 2013; Spagna et al., 2022), and use partially overlapping frontoparietal circuits (Behrmann et al., 2004; Landry et al., 2024; Shomstein, 2012; Spagna et al., 2022; Toba et al., 2023; Tosoni et al., 2023).

The Dorsal Attention Network (DAN), including the intraparietal sulcus, superior parietal lobule, and frontal eye fields, sends top-down signals supporting conscious report (Botta et al., 2014; Chica, et al., 2013; Corbetta et al., 2008; Dugué et al., 2020; Liu, et al., 2023). The predominantly right-lateralized Ventral Attention Network (VAN) supports target detection and reorienting (Corbetta et al., 2002; Doricchi et al., 2010; Liu, et al., 2023). These networks are interconnected by the three branches of the superior longitudinal fasciculus (SLF) (Thiebaut De Schotten et al., 2005; 2011): SLF I links regions within the DAN. SLF II connects angular gyrus and the middle frontal gyrus (MFG), bridging DAN and VAN (Fox et al., 2006; He et al., 2008); SLF III connects the supramarginal gyrus and the inferior frontal gyrus (IFG) within the VAN (Liu, et al., 2023; Rojkova et al., 2016). Disruption of SLF II/III impairs DAN-VAN interactions, with right hemisphere damage causing left visuospatial neglect (Bartolomeo et al., 2012; Thiebaut De Schotten et al., 2005; 2014; Doricchi et al., 2008; Lunven & Bartolomeo, 2017; Mengotti et al., 2020; Urbanski et al., 2011). An additional ventral temporo-occipital node has recently been identified within the network supporting endogenous attention (Sani et al., 2021); it exhibits attentional modulation comparable to dorsal frontoparietal cortices and maintains direct connectivity with frontoparietal regions. Characterizing oscillatory dynamics across attention nodes may clarify how neural communication prepares conscious perception (Cogitate Consortium et al., 2025; Mashour et al., 2020).

Here, we examined attention-driven oscillatory dynamics during the cue–target interval, to identify preparatory activity predicting target report or its omission. We focused on 18 anatomically defined regions of interest encompassing DAN and VAN nodes, their SLF-mediated connections, the temporo-occipital region, and bilateral visual cortices (Martín-Signes et al., 2024). Based on the dissociation between orienting and reorienting systems, we predicted that valid trials would preferentially engage DAN nodes via beta- and gamma-band oscillations and associated frontoparietal coherence reflecting top-down orienting (Buschman & Miller, 2007; Raposo et al., 2023; Xiong et al., 2023). In contrast, invalid trials would preferentially recruit VAN circuits through theta and alpha/beta dynamics consistent with beta- and theta–gamma-mediated coordination of right-lateralized ventral networks during reorienting (Fiebelkorn et al., 2018; Helfrich et al., 2018; Jensen et al., 2007; Proskovec et al., 2018), and with the pre-target “lookout” phenomenon (Liu et al., 2023).

## Method

### Participants

Of the 19 neurotypical participants that completed the experiment, five were excluded due to technical issues during acquisition. We analyzed magnetoencephalographic (MEG) data from *n* = 14 (7 female; 7 male), participants aged between 20–32 years (mean 24 ± 3.79 years) who took part in a previously published MEG study about the influence of spatial attention on the perception of near-threshold targets (Spagna et al., 2022). This study was promoted by the INSERM (C11–49) and approved by the Institutional Review Board of Île de France I. All participants were right-handed, had normal or corrected-to-normal vision, and gave written informed consent before participation.

### General Overview of the protocol

Participants arrived at the lab, provided written informed consent, and received instructions about the study. The study consisted of a threshold calibration session to determine individual contrast sensitivity, followed by a practice block and eight task blocks conducted in the MEG recording environment at the Paris Brain Institute - CENIR neuroimaging platform (https://parisbraininstitute.org/cenir-neuroimaging-platform). The experimental design, task procedure, and data were previously described in (Spagna et al., 2022); therefore, this study follows the same experimental procedure.

Participants were seated in a magnetically shielded room while stimuli were projected on a black projection screen positioned ∼80 cm away. The task was compiled and run using E-Prime software (RRID: SCR_009567; Psychology Software Tools, Pittsburgh, PA) on a Windows XP desktop computer, with a PROPixx projector (resolution: 1050 × 1400 pixels; refresh rate: 60 Hz) displaying the stimuli on a gray background.

### Experimental Design

#### Stimuli and Display

All stimuli were presented on a gray background at the center of a black projection screen using a PROPixx projector (resolution: 1050 × 1400 pixels; refresh rate: 60 Hz) located outside the shielded recording room. The display consisted of three rectangular placeholders, each one measuring 3.6◦ × 4.9◦ of visual angle. One was presented at the center of the screen and contained a black fixation cross. The other two placeholders were positioned at the bottom left and bottom right of the screen, each located 6° laterally and 4° below the central fixation cross. On each trial, a near-threshold Gabor patch target appeared in one of the peripheral placeholders. These stimuli were created with a maximum Michelson contrast of 0.92 (100%) and a minimum of 0.02 (1%). We chose to present stimuli in the lower quadrants to optimize MEG responses from early visual areas (Portin et al., 1999) (**Figure 1**).

**Figure 1.**
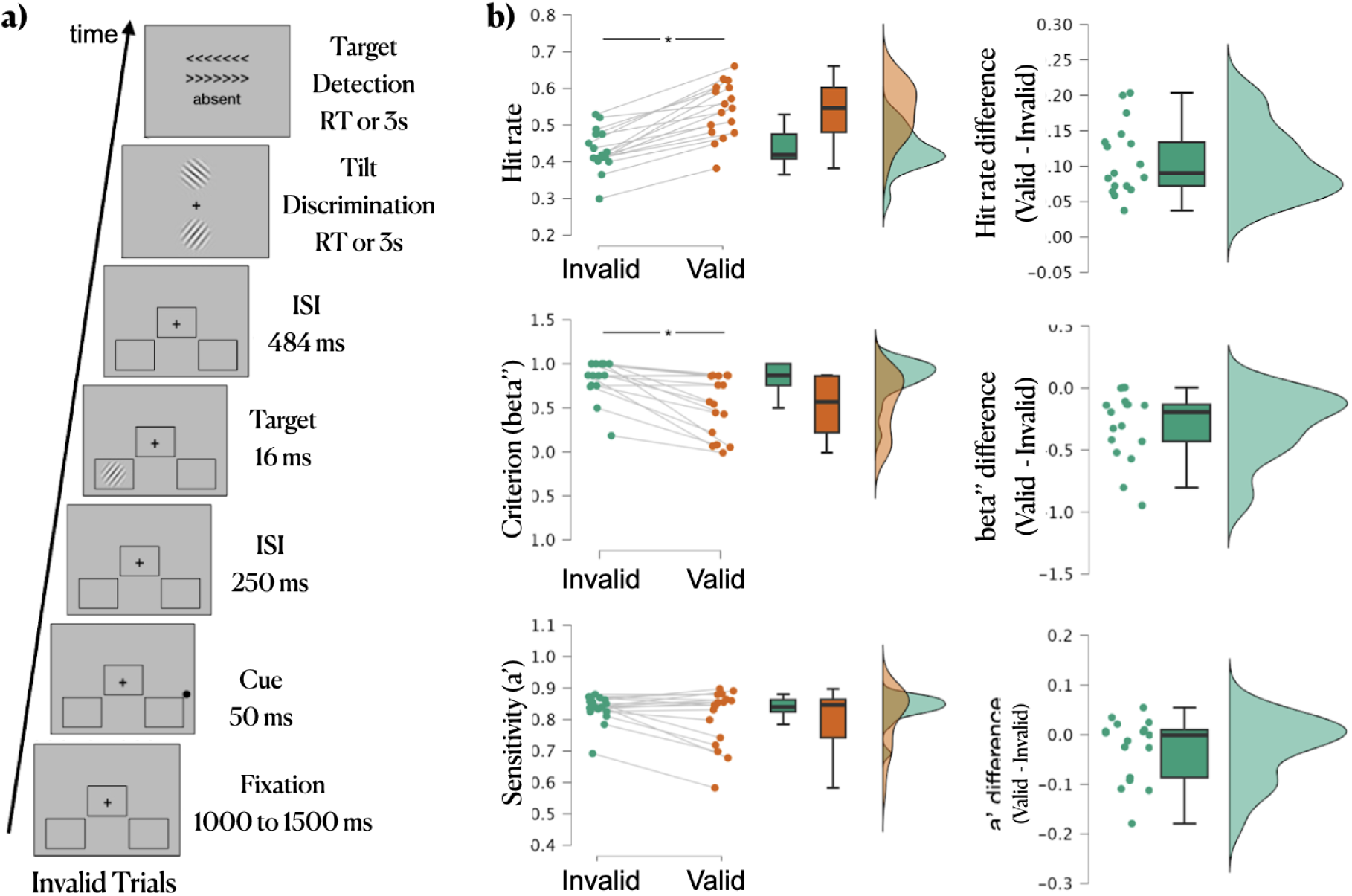
Task design and behavioral results. a) Stimulus Sequence: Schematic representation of the sequence of events for invalidly cued target trials. The display consisted of three rectangular placeholders: one at the center of the screen containing a black fixation cross and two at the bottom left and bottom right. On each trial, a near-threshold Gabor target appeared in one of the peripheral placeholders, preceded by a supra-threshold visual cue in the opposite location. In the tilt discrimination task, participants pressed a button to indicate the orientation of the Gabor patch. In the target detection task, participants indicated whether they did not see the target (“absent”), or whether it had been presented in the left (<<<<<<<) or right (>>>>>>>) target placeholder. b) Behavioral Performance: top panel: Hit rates were higher for validly than for invalidly cued targets (*p* < 0.001), as shown in the left raincloud plot, with the right plot illustrating the difference. Middle panel: the decision criterion (beta’’) was more liberal for validly cued targets (*p* < 0.01), indicating that valid cues increased the likelihood of reporting target presence. The left plot depicts the distribution of beta’’ values, while the right plot illustrates the difference between conditions. Bottom panel: perceptual sensitivity (a’) remained comparable across conditions (*p* = 0.14), suggesting that cue validity did not influence sensory evidence accumulation. The left plot shows the distribution of a’ values, while the right plot displays the difference between them. Additional analyses of these datasets (e.g., ANOVAs on accuracy) can be found in our previous study (Spagna et al., 2022).

#### Task Design

Each trial required participants to complete two sequential behavioral tasks: a tilt discrimination task, in which they judged the orientation of a Gabor patch, and a target detection task, in which they reported whether they had seen a target and, if so, in which visual hemifield it had appeared (**Figure 1a**). Each trial followed this sequence: (1) Fixation display (1000–1500 ms ISI): participants fixated on a central black cross. (2) Cue presentation (50 ms): a black dot (1° diameter, RGB: 128/128/128) appeared near one of the two peripheral placeholders. In 67% of trials (cued condition), the cue appeared near the placeholder where the target would later appear. In the remaining 33% of trials (uncued condition), the cue appeared in the opposite location. Participants were informed that the cue was predictive of the location of the target (i.e., in target-present trials, cues were likely to indicate the target location). (3) Inter-stimulus interval (ISI, 250 ms): a brief delay before target onset. (4) Target presentation (16 ms): a near-threshold Gabor patch (spatial frequency: 5 cycles/degree; diameter: 2.5°; orientation chosen among 12 equally spaced angles between 0° and 180°, excluding vertical and horizontal) appeared in one of the two peripheral placeholders. (5) Second ISI (484 ms): participants maintained fixation before the behavioral response phase began. (6) Tilt discrimination task: participants identified the orientation of the Gabor patch using a response box with three vertical buttons. Two reference orientations were displayed on the screen, differing by 3°. Participants pressed the upper button (index finger) for the upper orientation or the middle button (middle finger) for the lower orientation (only one of the two was correct, and the correct orientation was randomized across trials). The third button was not used in this task. After the response to the tilt discrimination task, or after 3 s of no response, (7) Target detection task. Two arrow-like stimuli (>>>>>> or <<<<<<) appeared above and below the fixation cross, with the word “absent” beneath them. Participants responded using the same three-button response box: upper button (index finger): Target seen in the right placeholder. middle button (middle finger): Target seen in the left placeholder. Lower button (ring finger): Target not seen. After a jittered 1–1.5 s interval, the next trial began.

#### Calibration Block

Prior to the main experiment, participants completed a calibration session (∼6 min) to determine the contrast threshold at which the target was detected ∼50% of the time. This session ensured that each participant’s contrast level was individually adjusted to yield approximately equal proportions of seen and unseen trials. A two-down/one-up adaptive staircase procedure psychophysical staircase procedure was used to estimate detection thresholds separately for validly and invalidly cued targets, thereby ensuring a balanced number of trials across the four main experimental conditions: seen valid (SV), seen invalid (SI), unseen valid (UV), and unseen invalid (UI). Participants performed the same task as described in the Task Design section, with the only difference being that the target contrast varied dynamically based on the participant’s prior responses.

#### Experimental Blocks

Following calibration, participants completed eight MEG recording blocks (∼8 min per block), for a total of 784 trials, distributed as follows: 448 validly cued target trials (67%), 224 invalidly cued target trials (33%), and 112 catch trials (no stimulus presented). The procedure is described in the Task Design section.

### MRI recordings

High-resolution T1-weighted structural MRI images (MPRAGE sequence, flip-angle, 9; Repetition Time, 2300 ms; Echo Time, 4.18 ms; voxel size: 1 x 1 x 1 mm) were acquired for each participant using a 3-T Siemens, TRIO whole-body MRI scanner (Siemens Medical Solutions, Erlangen, Germany) located at the CENIR MRI center (Paris Brain Institute France). After acquisition, images were segmented using the FreeSurfer “recon-all” pipeline (Fischl, 2012), and imported in Brainstorm (Tadel et al., 2011) for co-registration purposes. MEG sensors and structural MRI images were first manually aligned using the nasion/left ear/right ear (NAS/LPA/RPA) fiducial points recorded in the MEG file and in the MRI MNI coordinates. Co-registration was then refined using Brainstorm’s “Refine using head points” function, which applies an iterative closest point algorithm to fit the digitized head shape to the scalp surface. For additional methodological details (Tadel et al., 2019).

### MEG recordings

Continuous MEG recordings were conducted at the CENIR (https://parisbraininstitute.org/cenir-neuroimaging-platform) with an ELEKTA Neuromag TRIUX^®^ machine (204 planar gradiometers and 102 magnetometers) located in a magnetically shielded room. Data were recorded with a sampling frequency of 1,000 Hz and a bandwidth of 0.01–300 Hz. The recordings were then MaxFiltered (v2.2) (Taulu & Simola, 2006) to attenuate environmental noise. Signal Space Separation (SSS) was applied, bad channels were automatically detected, and data were band-pass filtered between 1–250 Hz, then resampled to 250 Hz. The resulting signals were converted to the FieldTrip structure (RRID: SCR_004849; http://www.fieldtriptoolbox.org/) (Oostenveld et al., 2011) for subsequent analysis in MATLAB. Cardiac activity (electrocardiogram – ECG), and vertical and horizontal Electrooculogram (EOG) signals were recorded to detect physiological artifacts. To ensure precise timing, stimulus onset was synchronized using a photodiode placed in the MEG room. The photodiode signal was used to exclude trials based on the following criteria: 1) trials with a delay between the trigger and the photodiode greater than 300 ms; 2) trials with a delay between the cue and the target greater than 827 ms; 3) trials in which the delay between the trigger of the cue and the photodiode was greater than 40 ms or smaller than 30 ms. Approximately 1% of trials were excluded based on these criteria.

#### Preprocessing and Artifact Rejection

Additional preprocessing steps were conducted using Fieldtrip (Oostenveld et al., 2011) with standard parameters. The process began with a visual inspection of continuous MEG data to identify and exclude segments containing artifacts or noise. Eye movement artifacts were rejected using EOG recordings from both vertical and horizontal sensors. Trials were excluded if eye movements exceeded (3° of visual angle). Rejection thresholds for both EOG traces were set to (±0.66 V), corresponding to the deviation beyond 3° (given that the target was presented at 6° of visual angle). Approximately 10.52% of trials were rejected offline from the MEG traces due to eye movements or blinks. In addition, trials contaminated by muscular artifacts (e.g., head or jaw movements) were manually rejected via visual inspection using FieldTrip’s ft_databrowser, amounting to ∼15% of total trials. Of the original 10,796 trials acquired, a total of 7,454 trials were retained for analysis. The final distribution was: *right visual field*: seen invalid = 396; seen valid = 1,588; unseen invalid = 812; unseen valid = 920; *left visual field*: seen invalid = 639; seen valid = 1,144; unseen invalid = 576; unseen valid = 1,379).

#### Event-Related Magnetic Fields

We analyzed data from the 102 magnetometer channels of the Elekta Neuromag TRIUX system. Although the TRIUX system records both magnetometers and planar gradiometers (102 and 204 channels, respectively), we restricted the present analyses to magnetometers. This decision was made for methodological consistency and computational efficiency in the context of experiment-wide permutation testing: magnetometers are more sensitive to deeper sources and produce broader spatial patterns, which are advantageous for visualizing bilateral and midline components. Planar gradiometers, while valuable, generate more focal maps and are generally more sensitive to superficial sources. Prior work using this system has demonstrated that analyses restricted to a single sensor type produce highly similar results to combined-sensor analyses (Wutz et al., 2014). Because all statistical inferences in the present study are based on source-reconstructed activity rather than sensor-level contrasts, restricting the forward model to magnetometers does not alter the fundamental signal content while substantially reducing computational demands.

Epochs were created using a Matlab (version: 9.13.0 (R2022b), The MathWorks Inc.) script, which segmented the continuous MEG recordings into 2,300 ms trials (from −1,000 ms before cue onset to +1,300 ms after the cue). Epochs were sorted into eight experimental conditions for each participant and imported into Brainstorm (Tadel et al., 2011). For each condition, event-related magnetic fields were computed by averaging all retained trials across the entire epoch duration (−1000 to +1300 ms relative to cue onset; total length = 2300 ms).

#### Source reconstruction

Signal amplitudes from the 15,002 cortical elemental dipoles underlying the signals measured by the sensors were then estimated from the epochs using the weighted minimum norm estimation (wMNE) imaging method as implemented in Brainstorm (Tadel et al., 2011). The number of dipoles used in our study is a standard in MEG literature, and is meant to cover the entire cortical surface and its folded anatomy (Tadel et al., 2019). The wMNE method identifies a current source density image fitting the data through the forward model, and then favors solutions that are of minimum energy by using source covariance as a prior. wMNE modeling of the sources was used because it is considered to be the gold standard in the field, because it balances between a conservative approach in reconstructing sources of MEG signal, with reliable statistical inference across participants, and being computationally efficient (less resource-expensive) (Baillet et al., 2001; Tadel et al., 2019). Subject-specific noise covariance matrices were computed from the pre-cue baseline window (−1000 to −2 ms). Estimating the baseline covariance (i.e, noise covariance) over such an amount of time (approximating one second) allows us to assume a sufficient regularization term of the solution (noise from brain activity at rest thought to be unrelated to our task conditions). A constrained source covariance model was used to model one dipole, oriented normally to the surface, a choice that allows us to study the cortical regions associated with the interaction between attention and consciousness. No smoothing kernel was applied, and sources were then re-interpolated (projected) on a common template (MNI 152).

#### Regions of Interest (ROIs)

We adopted an ROI-based approach for three key reasons. First, our primary focus was to characterize the neural dynamics underlying conscious visual perception of attended and unattended targets. To do so, we included sensory regions together with frontoparietal cortices, allowing us to comprehensively examine both local activity and inter-regional communication within networks known to support visual information processing (Fiebelkorn et al., 2018; Fries, 2023; Helfrich et al., 2018; Liu, Bayle, et al., 2023; Spadone et al., 2021; Spagna et al., 2022; Tosoni et al., 2023; Xiong et al., 2023). In addition, our set of ROIs encompassed a ventrotemporal attention node and two nodes in the occipital cortex (one lateral and one medial) to corroborate the existence of distinct oscillation patterns during seen and unseen trials in low-and mid-level visual regions, a topic that is still being explored (Dugué et al., 2020; Esghaei et al., 2022a; Tosoni et al., 2023). Second, frontoparietal regions are consistently implicated in attention and conscious perception (Bartolomeo et al., 2025; Geng et al., 2019; Helfrich et al., 2018; Liu, Bayle, et al., 2023; Spagna et al., 2022; Szczepanski et al., 2014), making them highly relevant for investigating predictive processing and target detection mechanisms. Third, restricting our analyses to predefined ROIs increases statistical power and interpretability. This approach enables testing of specific hypotheses about functional roles while minimizing the risks associated with whole-brain exploratory analyses and multiple comparison corrections.

Eighteen ROIs were defined anatomically in both right and left hemispheres, using the Scout function in Brainstorm (unconstrained mode). Each ROI was created by identifying a seed vertex within a Brodmann area (BA) and then manually expanding the scout to approximately cover the full extent of the corresponding BA. All ROIs were standardized to include 120 vertices, corresponding to an average area of approximately 16 cm². **Supplementary Table 1** reports the seed vertex index, MNI coordinates, corresponding BA, and surface area for each ROI.

Six regions were drawn along the boundaries of frontal and parietal projections of the dorsal, intermediate, and ventral branches of the superior longitudinal fasciculus (respectively, SLF I, II, and III). An atlas detailing frontoparietal connections along the SLFs was used to reference which BA to include within each ROI (Rojkova et al., 2016). For the regions pertaining to the SLF I, an ROI in the superior frontal gyrus (SFG) was defined as the region within BA8 and BA9 bounded ventrally by the superior frontal sulcus, posteriorly by BA6 and anteriorly by BA10. The parietal projection of the SLF I was defined as the portion of BA7 bounded between the superior parietal sulcus (SPL), posteriorly by BA19 and anteriorly by BA40. For the regions pertaining to the SLF II, an ROI in the middle frontal gyrus (MFG) was defined as a region within BA46 and bounded dorsally by the superior frontal sulcus, ventrally by the inferior frontal sulcus of the pars opercularis (BA44) and pars orbitalis (BA47), anteriorly by BA10 and posteriorly by the primary motor cortex (BA4). The parietal projection of the SLF II was defined as the angular gyrus (BA39) (IPS). For the SLF III projections, an ROI in the inferior frontal gyrus (IFG) was defined as encompassing the inferior frontal sulcus from the pars opercularis (BA44) to the pars orbitalis (BA47), bounded anteriorly by BA10 and posteriorly by the primary motor cortex (BA6). The parietal projection of the SLF II was defined as the supramarginal gyrus (BA40) (TPJ). Another ROI was drawn along a previously identified ventro-temporal cortical node (TO node) (BA37) associated with the attentional control network (MacLean et al., 2023; Sani et al., 2021). Two further ROIs targeted early visual cortex: one along the lateral occipital surface encompassing V1 and V2 (BA18), and another along the medial occipital surface on the calcarine sulcus (BA17) to capture attentional effects on peripheral and foveal regions of the visual cortex. See **Figure 2** and **Supplementary Table 1** for more details about the area included in each ROI, including centroids in MNI coordinates and extents.

**Figure 2.**
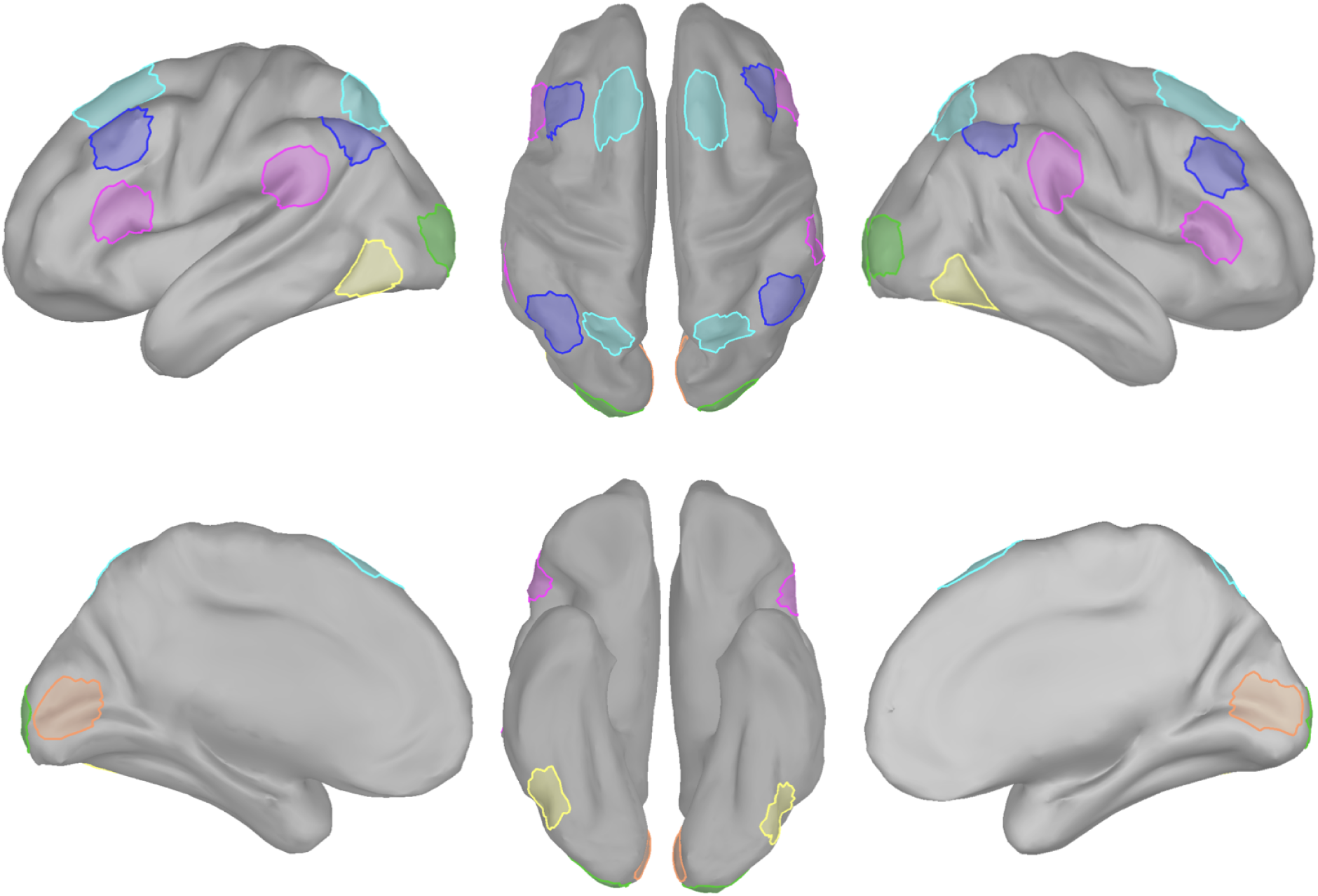
Surface area for each of the 18 regions of interest. The figure displays regions linked by SLF I (teal), SLF II (blue) and SLF III (pink). Yellow: ventro-temporal cortical node. Orange: medial visual cortex. Green: lateral visual cortex.

## Data Analysis

### Behavioral performance

Signal detection theory (SDT) was used to assess perceptual sensitivity (a’) and response bias (beta’’) based on the spatial relationship between cue and target (valid vs. invalidly cued targets). A’ quantifies the signal-to-noise ratio, where values range from 0.5 (chance-level) to 1 (perfect performance), while beta’’ reflects response criterion, with values near 1 indicating a conservative approach (fewer false alarms, more misses) and values near −1 indicating a liberal approach (more false alarms).

Target contrast was calibrated to achieve ∼50% detection at both valid and invalid locations. Given that some participants made no false alarms, nonparametric SDT measures were used (Pollack & Norman, 1964). We computed the percentage of seen targets (hits) and false alarms (FAs), estimating a’ and beta’’ separately for valid and invalid trials. Trials were partitioned in seen or unseen based on participants’ response to the detection task: seen trials comprised only trials when the participant also reported the correct left/right position of the target (Chica et al., 2010). This criterion ensured high confidence in the classification of trials as consciously perceived.

Differences between conditions were tested using paired-sample t-tests, while Wilcoxon signed-rank tests assessed potential differences in threshold sampling during calibration. Additional analyses of these datasets (e.g., ANOVAs on accuracy) can be found in our previous study (Spagna et al., 2022).

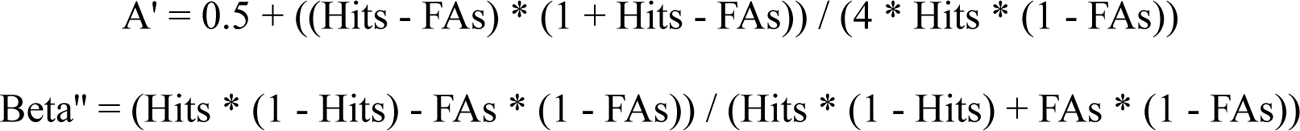

### Oscillatory activity within each ROI

Time–frequency decomposition of source-level activity was performed using complex Morlet wavelets in Brainstorm (Tadel et al., 2011), with a central frequency factor (MorletFc) of 1 and temporal resolution (MorletFwhmTc) of 3. The method computed power values across 67 frequencies ranging from 4 to 70 Hz, at 501 time points spanning −500 to 500 ms relative to cue onset. This analysis was performed on PCA-extracted scout time series (function: pca2023) from 18 anatomically defined ROIs (9 per hemisphere), each reduced to a single representative component via PCA. To isolate cue-related changes, baseline correction was performed using z-score normalization relative to the −500 to −100 ms pre-cue window, as implemented in Brainstorm’s process_baseline_norm. The resulting time–frequency maps reflect baseline-normalized gamma power in units of standard deviation (z).

To identify condition-related changes in oscillatory activity, we conducted a non-parametric one-sample t-test using a max-T permutation procedure (FieldTrip, ft_freqstatistics; 10,000 permutations; significance threshold of α = 0.05) on within-subject contrast maps derived from time–frequency decompositions. For each of the 18 ROIs, average power was computed for the four experimental conditions: Seen Valid (SV), Seen Invalid (SI), Unseen Valid (UV), and Unseen Invalid (UI), focusing on the 4–70 Hz frequency range and a post-cue window from −100 ms to 400 ms. Two contrasts were estimated for each participant:

1. the main effect of Report = ((SV + SI)/2 − (UV + UI)/2)
2. the Report × Validity interaction = ((SI − SV) − (UI − UV)).

Multiple comparisons were controlled across the full ROI × frequency × time grid, ensuring that the correction accounted for the entire spatiotemporal-spectral search space. No spatial adjacency constraints were imposed since the analysis was ROI-based. This procedure identified individual time–frequency tiles within each ROI where the contrast significantly differed from zero across participants. For each significant tile, we extracted the corrected *p*-value, peak *t*-value, spectral and temporal location, and within-subject effect size (Cohen’s *d*z), obtained from the comparison of paired conditions within subjects. To further interpret significant interaction effects, follow-up pairwise comparisons were conducted using within-subject simple effects analyses. Specifically, we compared SI versus SV and UI versus UV conditions within each significant ROI to determine whether the interaction was driven by differences in the validity effect between Seen or Unseen trials. These targeted contrasts clarified the contribution of attentional validity to conscious report, and confirmed that interaction effects were supported by distinct patterns across cueing conditions.

### Coherence among ROIs

To characterize inter-regional coordination during anticipatory attention, we computed imaginary coherence between all pairs of predefined ROIs across five canonical frequency bands: theta (4–7 Hz), alpha (8–13 Hz), beta (14–29 Hz), low gamma (30–59 Hz), and high gamma (60–90 Hz). Imaginary coherence isolates the interaction component not explainable by zero-lag correlations, thereby reducing the influence of volume conduction and field spread (Nolte et al., 2004). Coherence was estimated using Brainstorm’s process_cohere1n (Hilbert-based connectivity), which indexes the linear relationship between complex-valued analytic signals in the frequency domain (Brookes et al., 2011; Xiong et al., 2023). ROIs were anatomically defined (see Methods), and signals were extracted using PCA (pca2023 function in Brainstorm). All data were *z*-score normalized relative to a common pre-cue baseline (–500 ms to –100 ms) before coherence computation. At the group level, we tested two within-subject contrasts using a non-parametric max-T permutation framework (FieldTrip, ft_freqstatistics; 10,000 permutations; significance threshold of α = 0.05):

1. the main effect of Report ((SV + SI)/2 − (UV + UI)/2) and
2. the Report × Validity interaction ((SI − SV) − (UI − UV)).

For each ROI pair, frequency band, and time point, difference scores were submitted to a one-sample *t*-test against zero, and the maximum *t*-value across the full ROI × frequency × time space was used to build the null distribution, providing control of the family-wise error rate at α = .05. Significant interaction effects were further examined using pairwise simple effect analyses (SI vs. SV and UI vs. UV) within each significant ROI to determine whether the interaction was driven by differences in the validity effect between Seen or Unseen trials. For each significant ROI pair, we report the corrected *p*-value, peak *t*-value, frequency band and temporal location of the effect, and the corresponding within-subject effect size (Cohen’s *d*), obtained from the comparison of paired conditions within subjects.

### Phase-Amplitude Coupling within each ROI

To examine whether participants’ attentional state modulated cross-frequency coupling, we computed theta–gamma phase-amplitude coupling (PAC) using the pipeline implemented in Brainstorm (Canolty & Knight, 2010; Voytek, 2010). PAC estimation was performed separately for each participant, ROI (n = 18), and task condition (Seen Valid, Seen Invalid, Unseen Valid, Unseen Invalid).

PAC was estimated over the cue–target interval (0–300 ms post-cue) using the direct PAC measure (sPAC.DirectPAC) in Brainstorm. All analyses were applied to the scout time series extracted from anatomically defined ROIs using PCA decomposition (function: pca2023). This step ensured that the PAC computations were based on low-dimensional, source-level representations, and reduced computational demands. To allow consistent interpretation of neural dynamics across conditions, PAC values were normalized using z-score transformation relative to a common pre-cue baseline interval (–500 to –100 ms relative to cue onset). This baseline correction was applied prior to group-level analysis and ensures comparability across ROIs, conditions, and frequency bins. For each ROI, raw time series were band-pass filtered in the theta (4–8 Hz) and gamma (30–70 Hz) ranges. Instantaneous phase and amplitude were extracted using the analytic signal. PAC was computed as the modulation index between phase (five frequencies from 4 to ∼8 Hz) and amplitude (17 gamma frequencies from 30 to 70 Hz), resulting in 5 × 17 PAC matrices per condition and ROI. These were averaged across trials to obtain a single PAC map per participant, condition, and ROI. Output values were saved in .mat files for group-level analysis.

At the group level, we submitted the 5 × 17 PAC matrices to a within-subjects statistical comparison using a non-parametric max-T test (FieldTrip, ft_freqstatistics; 10,000 permutations; significance threshold of α = 0.05) for the following contrasts:

1. the main effect of Report ((SV + SI)/2 − (UV + UI)/2)
2. the Report × Validity interaction ((SI − SV) − (UI − UV)).

Statistical testing was performed per ROI, correcting for multiple comparisons across the full PAC grid (phase × amplitude frequency) using the maximum t-value across all bins as the null distribution. Only bins surviving this correction (FWER ≤ 0.05) were interpreted. For each ROI showing at least one significant bin, we report the corrected *p*-value, the phase and amplitude frequency ranges containing significant bins, and the mean Cohen’s d obtained from the comparison of paired conditions within subjects. In addition, significant interaction effects were followed up with pairwise simple effect analyses (SI vs. SV and UI vs. UV) within each significant ROI to determine whether the interaction was driven by differences in the validity effect between Seen or Unseen trials.

## Results

### Decision criterion shifts for validly cued targets

Hit rates were higher for validly cued targets (M = 0.53, SD = 0.07) than for invalidly cued targets (M = 0.43, SD = 0.06), (*t* = 7.57, *p* < 0.01, *d* = 2.02). Perceptual sensitivity (a’) remained comparable for valid (M = 0.82, SD = 0.08) and invalidly cued targets (M = 0.84, SD = 0.03), (*t* = 1.57, *p* = 0.14, *d* = 0.14), suggesting that cue validity did not affect sensory evidence accumulation. However, participants adopted a more liberal criterion for validly cued targets (M = 0.56, SD = 0.34) compared to invalidly cued targets (M = 0.88, SD = 0.15), (*t* = 3.99, *p* < 0.01, *d* = 1.07), suggesting that cues biased decision-making and participants were more likely to report target presence after valid cues (**Figure 1**).

### Beta oscillations in right-lateralized attentional nodes reflects the interaction between attention and conscious reports

The contrast of Report × Validity revealed an interaction in the right superior parietal lobule (SPL), where beta-band activity (∼26 Hz) was modulated by the combination of cue validity and perceptual report (**Figure 3, panels a–d**). This effect emerged early in the cue–target interval (58 ms), with a peak t-value of –5.89 and a large within-subject effect size (Cohen’s d = –1.57). Specifically, beta amplitude was lower in the Seen Invalid (SI) condition (mean = 0.43, SD = 0.20) compared to the Seen Valid (SV) condition (mean = 0.59, SD = 0.12), confirmed by a significant simple effect (*t*(13) = –2.36, *p* = 0.03, *d* = –0.63). A crossover pattern was observed in the Unseen conditions, with beta amplitude in the Unseen Valid (UV) condition (mean = 0.47, SD = 0.15) being lower than in the Unseen Invalid (UI) condition (mean = 0.58, SD = 0.21); however, this simple effect did not reach significance (*t* = 1.99, *p* = 0.07, *d* = 0.53). This suggests that early SPL beta oscillations support sustained attention at the cued location. When the cue is valid, increased beta amplitude early in the cue–target interval appears to support exogenous attentional capture, facilitating conscious perception. Conversely, when the cue is invalid, this mechanism may impede the required reorienting toward the uncued target, leading to more perceptual misses.

**Figure 3.**
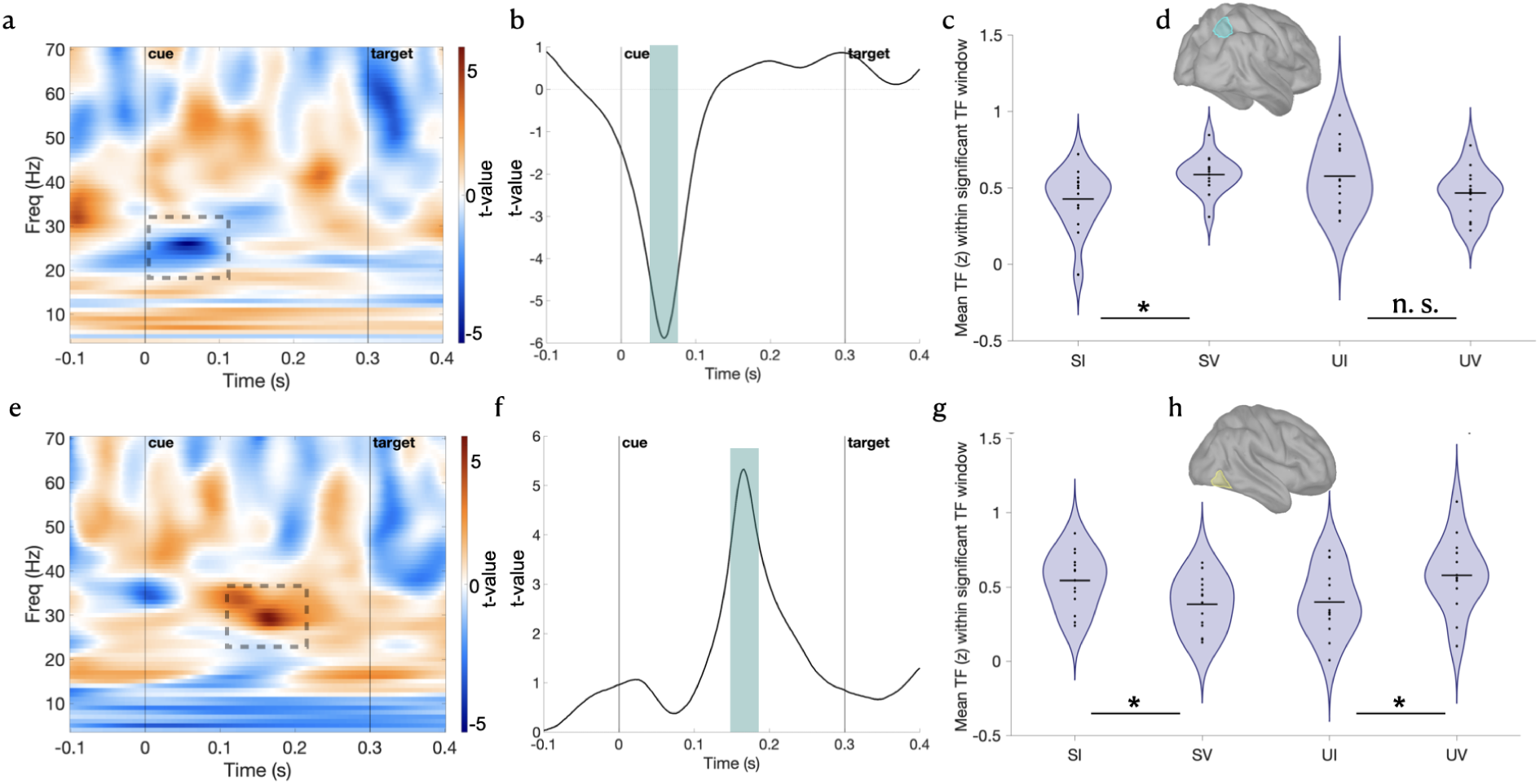
Beta oscillations in right-lateralized attentional nodes reflect the interaction between attention and conscious reports. *Top row:* results for the right Superior Parietal Lobe. **(a)** Time–frequency map (*t*-values) showing a significant interaction in the beta band (∼26 Hz) emerging at ∼58 ms post-cue. **(b)** Temporal evolution of *t*-values at 26 Hz. The shaded area indicates the time window of the significant effect. **(c)** Violin plot of condition means reveals a crossover interaction: SI = 0.43 (SD = 0.20); SV = 0.59 (SD = 0.12); UI = 0.58 (SD = 0.21); UV = 0.47 (SD = 0.15). **(d)** teal mask contours the right SPL. This pattern suggests that SPL beta activity tracks the interaction between perceptual outcome and cue validity (Cohen’s d = −1.57). *Bottom row:* results for the right Temporo-Occipital Node. **(e)** Time–frequency map (*t*-values) showing a significant interaction in the beta band (∼29 Hz) emerging at 166 ms post-cue. **(f)** Temporal evolution of *t*-values at ∼29 Hz; shaded area marks the significant effect. The shaded area indicates the time window of the significant effect. **(g)** Violin plot of condition reveals a crossover interaction:SI = 0.54 (SD = 0.19); SV = 0.39 (SD = 0.18); UI = 0.40 (SD = 0.22); UV = 0.58 (SD = 0.25). This pattern suggests that beta activity in the RH TO node flexibly tracks perceptual success and cue expectation. **(h)** yellow mask contours the right TO ROI. An asterisk indicates statistically significant simple effects (max-T permutation, 10,000 iterations, two-tailed, FWER .05; *p* < .05).

A second interaction emerged in the right temporo-occipital (TO) node, where beta-band activity (∼29 Hz; **Figure 3, panels e–h**) was modulated by the combination of perceptual report and cue validity. This interaction emerged at 166 ms post-cue, with a peak *t*-value of 5.33 and a large within-subject effect size (Cohen’s *d* = 1.42). Beta amplitude was greater in the Seen Invalid (SI) condition (mean = 0.54, SD = 0.19) compared to the Seen Valid (SV) condition (mean = 0.39, SD = 0.18), confirmed by a simple effect (*t* = 2.65, *p* = 0.02, *d* = 0.71). Conversely, amplitude was lower in the Unseen Invalid (UI) condition (mean = 0.40, SD = 0.22) compared to the Unseen Valid (UV) condition (mean = 0.58, SD = 0.25; *t*(13) = –2.24, *p* = 0.04, *d* = –0.60). This pattern suggests that β oscillations in the right TO node reflect preparatory reorienting processes that can either facilitate or hinder conscious perception depending on cue validity. Higher β before target onset in invalid trials (SI > SV) indicates that increased preparatory reorienting successfully compensated for the misleading cue, enabling detection of uncued targets. In contrast, elevated β in UV trials—where the cue was valid but the target was missed—implies that the same reorienting tendency became detrimental when attention should have remained at the cued location. This pattern supports a role for right TO beta oscillations in voluntary shifts of attention during the cue–target interval, shaping the interaction of endogenous attention and visual awareness.

In addition to these interactions, two main effects of Report occurred. First, in the left medial visual cortex, gamma-band activity (∼61 Hz) was reduced for Seen (mean = 0.11, SD = 0.07) compared to Unseen trials (mean = 0.20, SD = 0.07) **Supplementary Figure 1, panels a-c)**. This effect emerged at 104 ms post-cue, with a peak *t*-value of −5.71 and a large within-subject effect size (Cohen’s *d* = −1.53), suggesting that early gamma suppression in this region may index successful perceptual processing. Second, a complementary effect emerged in the left middle frontal gyrus (MFG), where high gamma activity (∼70 Hz) was enhanced for Seen trials (mean = 0.23, SD = 0.07) relative to Unseen trials (mean = 0.13, SD = 0.06) (**Supplementary Figure 1, panels d-f)**. This effect peaked later, at 282 ms, with a peak *t*-value of −5.72 and exhibited a large effect size (Cohen’s *d* = 1.53), suggesting that left-lateralized gamma oscillations in the MFG support perceptual processing of upcoming targets.

Together, these results reveal a distributed, frequency-specific pattern of beta and gamma activity across right temporo-occipital, right parietal, and left frontal regions. Beta oscillations in right-lateralized regions—including the superior parietal lobule and temporo-occipital cortex—were associated with successful conscious perception across both early and late stages of the cue–target interval. In contrast, left-lateralized high gamma activity supported conscious reports in late (i.e., middle frontal gyrus), but not early (i.e., medial visual region) stages of the cue-target interval.

### Interaction Between Attention and Awareness Modulates Interhemispheric Gamma-Band Coherence

To assess whether cross-regional oscillatory interactions contribute to conscious perception, we examined coherence between ROIs using a non-parametric max-T test. An interaction emerged in the high gamma band between the Right Middle Frontal Gyrus (MFG) and the Right Temporo-Occipital (TO) Node, peaking around 66 ms post-stimulus (*t* = –4.77, *p* < 0.0001, Cohen’s *d* = –1.27; **Figure 4, panels a-d**). This crossover pattern indicates that high fronto-temporal coherence in the gamma range early during the cue-target period disrupts conscious processing of invalidly cued targets. Simple effects confirmed this interaction: coherence was higher for UI (mean = 0.08, SD = 0.04) vs. UV (mean = 0.04, SD = 0.03; *t* = 3.36, *p* = 0.005, *d* = 0.90), but not for SI (mean = 0.04, SD = 0.04) vs. SV (mean = 0.05, SD = 0.05; *t* = –0.90, *p* = 0.39, *d* = –0.24).

**Figure 4.**
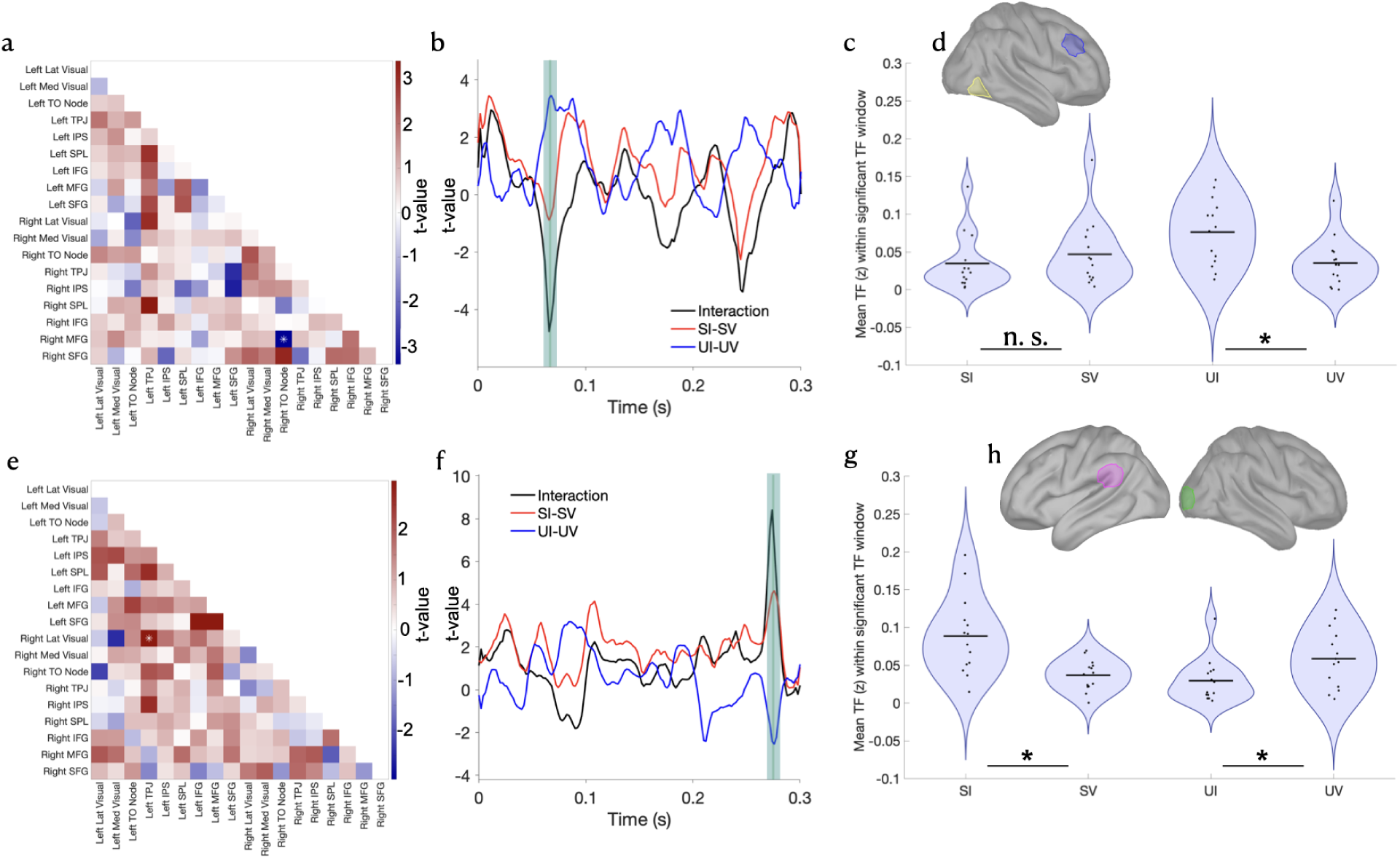
Report × Validity interactions in gamma-band coherence. *Top row* (a–d): A non-parametric max-T test revealed a significant interaction in the high gamma band (∼66 ms post-stimulus) between the Right Temporo-Occipital (TO) Node and the Right Middle Frontal Gyrus (MFG). The connectivity matrix **(a)** shows *t*-values, with a white asterisk marking the significant connection. The full list of ROI names and order is provided in the Supplementary Table 1. **(b)** Temporal evolution of *t*-values at the significant connection. The shaded area indicates the time window of the significant effect. Violin plots **(c)** display mean *z*-transformed coherence values, showing higher coherence in the Unseen Invalid (UI) compared to Unseen Valid condition, while the difference between Seen Invalid and Seen Valid was not significant. This interaction suggests increased fronto-temporal communication under mismatched expectations. Inset **(d)** contour of the cortical mask of the ROIs involved in this significant connection. *Bottom row* (e–h): A second significant interaction was observed in the low gamma band (∼274 ms) between the Right Lateral Visual cortex and the Left Temporo-Parietal Junction (TPJ). The connectivity matrix **(e)** highlights the *t*-values, with a white asterisk marking the significant connection. The full list of ROI names and order is provided in the Supplementary Table 1. **(f)** Temporal evolution of *t*-values at the significant connection. The shaded area indicates the time window of the significant effect. Violin plots **(g)** show higher coherence in the Seen Invalid (SI) condition compared to the Seen Valid condition, and higher coherence in the Unseen Valid condition (UV) compared to the Unseen Invalid (UI) condition. This crossover pattern suggests that low gamma connectivity between early visual and the left temporo-parietal region may support compensatory processing during violations of spatial expectations. Inset **(h)** shows the cortical mask of the ROIs involved in this significant connection. An asterisk indicates statistically significant simple effects (max-T permutation, 10,000 iterations, two-tailed, FWER .05; *p* < .05).

A second interaction occurred in the low gamma band between the Right Lateral Visual cortex and the Left Temporo-Parietal Junction (TPJ), emerging around 274 ms post-stimulus (*t* = 8.41, *p* < 0.0001, Cohen’s d = 2.25; **Figure 4, panels e-h**), indicating higher coherence for the Seen Invalid (mean = 0.09, SD = 0.05) condition compared to Seen Valid (mean = 0.04, SD = 0.02; *t* = 4.50, *p* = 0.0006, *d* = 1.20) and for Unseen Invalid (mean = 0.03, SD = 0.03), compared to Unseen Valid (mean = 0.06, SD = 0.04; *t* = –2.42, *p* = 0.03, *d* = –0.65). This result points to a later-occurring, low gamma-range increase in cross-hemispheric connectivity preceding successful perceptual processing of invalidly cued targets.

In addition to the two interaction effects, two main effects of Report revealed significant coherence modulation in both beta and gamma frequency bands (**Supplementary Figure 2**). An early effect (∼ 102 ms) in the beta-band emerged between the Right Superior Frontal Gyrus (SFG) and the Left TO Node (*t* = –6.74, *p* < 0.0001, Cohen’s *d* = −1.80), indicating greater coherence for Unseen trials than (mean = 0.08, SD = 0.03) than in the Seen condition (mean = 0.04, SD = 0.02). This suggests that fronto-temporal beta-band coupling is enhanced when stimuli fail to reach conscious awareness. A second Report-related effect was identified in the low gamma band (∼216 ms) between the Right MFG and the Left Inferior Frontal Gyrus (IFG). This effect (*t* = 7.65, *p* < 0.0001, Cohen’s *d* = 2.05) indicated greater coherence for Seen (mean = 0.08, SD = 0.02) compared to Unseen (mean = 0.04, SD = 0.02). Thus, enhanced cross-hemispheric prefrontal communication in the gamma band supports the conscious perception of upcoming targets.

Overall, these results suggest a frequency-specific dissociation between conscious and unconscious processing. Increased beta or high gamma coherence during the early cue–target interval led to more misses, whereas cross-hemispheric low gamma coherence was associated with better conscious reports of upcoming targets.

### Theta–Gamma cross-frequency coupling in the left inferior frontal gyrus is associated with conscious report

To investigate whether cross-frequency coupling contributes to conscious perception, we examined phase–amplitude coupling (PAC) across phase (4–8 Hz) and amplitude (30–70 Hz) frequencies. A non-parametric max-T test identified a significant Report × Validity interaction at the PAC combination of 7 Hz (theta phase) and 36 Hz (gamma amplitude) in the right temporo-occipital (TO) node (**Figure 5, panels a–e**). This effect was driven by elevated PAC strength in the Unseen Invalid (UI) condition (mean = 0.126, SD = 0.0025), relative to Seen Invalid (SI; mean = 0.121, SD = 0.0023), Seen Valid (SV; mean = 0.121, SD = 0.0022), and Unseen Valid (UV; mean = 0.120, SD = 0.0026), as shown in Figure 4b–c. This pattern suggests that enhanced theta–gamma coupling in the TO node may disrupt perceptual processing of targets preceded by invalid cues, when attentional shifts are required. However, follow-up simple effects did not reach statistical significance (SI vs. SV mean difference = 0.001, *t* = 1.78, *p* = 0.10, *d* = 0.4; UI vs UV *p* > 0.3) because PAC differences between conditions were small, with no reliable validity effect in either the Seen condition (SI vs. SV) or the Unseen condition (UI vs. UV).

**Figure 5.**
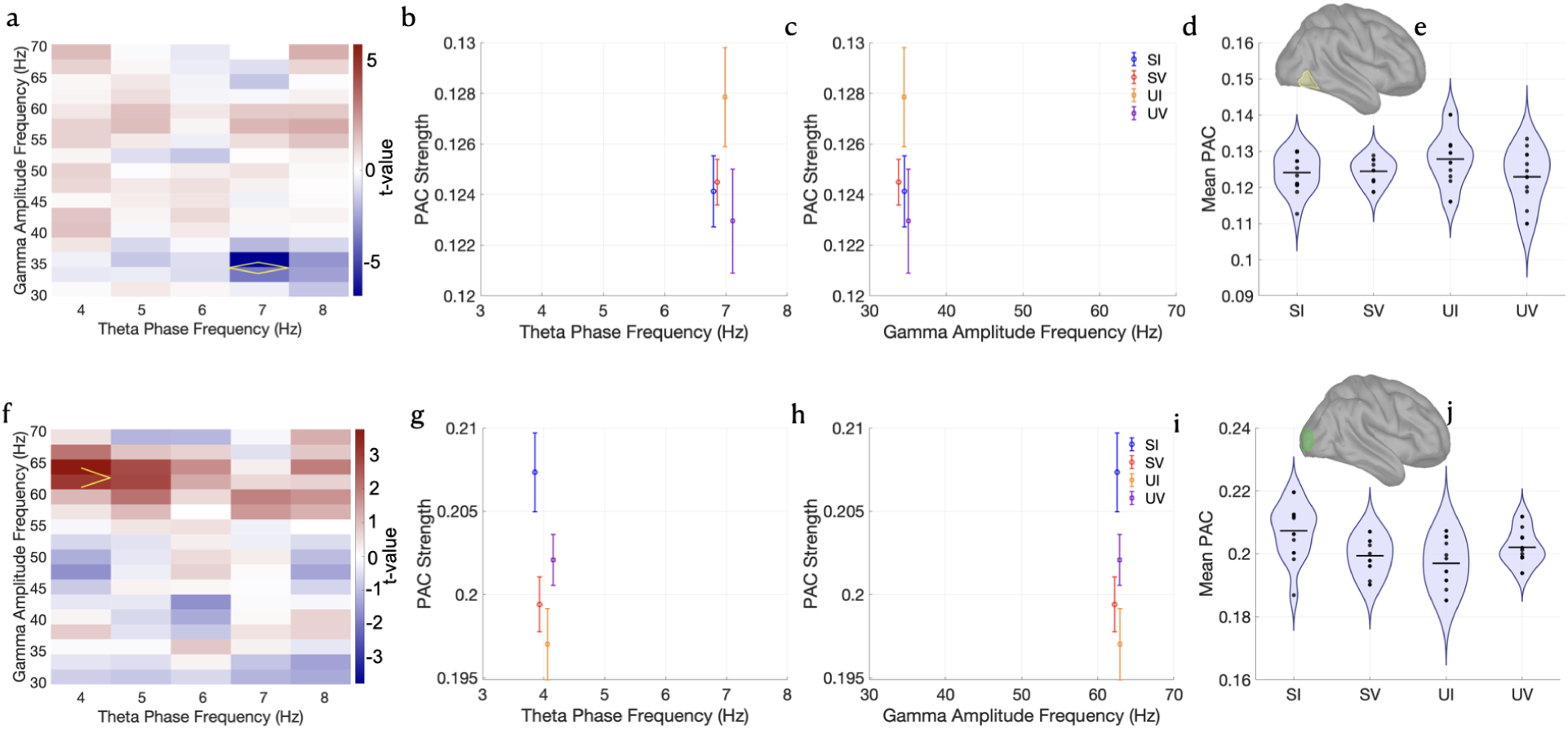
Top row: Interaction effect (Report × Validity) in phase–amplitude coupling (PAC) at the right temporo-occipital (TO) node (max-T permutation, 10,000 iterations, two-tailed, FWER .05; *p* < .05). **(a)** T-value matrix across phase (x-axis) and amplitude (y-axis) frequencies. A non-parametric max-T test identified a significant interaction effect at the combination of 7 Hz (theta phase) and 36 Hz (gamma amplitude), indicated by the yellow contour (*p* < .001). **(b, c)** PAC strength as a function of (**b**) theta phase frequency (x-axis) and **(c)** gamma amplitude frequency (x-axis) across conditions. The UI condition (orange) shows the highest PAC strength at 7 Hz (mean = 0.126, SD = 0.002), relative to SI (mean = 0.121, SD = 0.002), SV (mean = 0.122, SD = 0.002), and UV (mean = 0.121, SD = 0.002). Error bars denote ±1 standard deviation. **(d)** Violin plots depict the distribution of PAC strength within the significant time–frequency window across subjects for each condition, illustrating the interaction pattern and inter-individual variability. **(e)** the yellow mask contours the right TO node ROI. Bottom row: Interaction effect (Report × Validity) in phase–amplitude coupling (PAC) at the Right lateral visual cortex. **(f)** A significant interaction was detected at 4 Hz (phase) and 63 Hz (gamma amplitude). **(g, h)** PAC strength as a function of **(g)** theta phase frequency (x-axis) and **(h)** gamma amplitude frequency (x-axis) across conditions. **(i)** Violin plots showing PAC strength followed the pattern SI (mean = 0.207, SD = 0.009), UV (mean = 0.201, SD = 0.005), SV (mean = 0.199, SD = 0.006), UI (mean = 0.197, SD = 0.007). **(j)** the green mask contours the right lateral visual cortex ROI.

A second Report × Validity interaction was observed in the right lateral visual cortex at a PAC combination of 4 Hz (theta phase) and ∼63 Hz (gamma amplitude) (**Figure 5, panels f–j**). PAC strength was highest in the Seen Invalid (SI) condition (mean = 0.207, SD = 0.009), followed by Unseen Valid (UV; 0.201, SD = 0.005), Seen Valid (SV; 0.199, SD = 0.006), and Unseen Invalid (UI; 0.197, SD = 0.007), yielding a positive interaction contrast. This pattern indicated that increased theta–gamma coupling in this region may contribute to redirecting attention away from the expected (but incorrect) location, allowing for successful target detection at the uncued site. However, follow-up simple effects did not reach statistical significance (SI vs. SV: *t* = 1.12, *p* = 0.29; all other contrasts *p* > 0.5), because PAC differences between conditions were small, with no reliable validity effect in either the Seen condition (SI vs. SV) or the Unseen condition (UI vs. UV).

Two additional PAC effects reached significance across distinct cortical regions. In the left TO node, PAC strength at ∼6 Hz (theta phase) and ∼42 Hz (gamma amplitude) showed a significant main effect of Report (**Supplementary Figure 3, a–d**), with PAC being elevated for Unseen trials (mean = 0.183, SD = 0.005) compared to Seen trials (mean = 0.179, SD = 0.006), reflecting a negative contrast. This suggests that increased theta–gamma coupling in left TO leads to unsuccessful perceptual processing.

A second main effect of Report was observed in the left inferior frontal gyrus (IFG), at ∼7 Hz (theta) and 40 Hz (gamma) (**Supplementary Figure 3, e–h**). Here, PAC was higher for Seen trials (mean = 0.140, SD = 0.007) relative to Unseen trials (mean = 0.135, SD = 0.007). This suggests that theta–gamma coupling in the IFG scales with conscious perception regardless of cue validity, potentially reflecting the involvement of frontal networks in supporting conscious reports.

## Discussion

Neuronal oscillations, shaped by excitatory–inhibitory dynamics, organize into frequency-specific rhythms that synchronize across brain regions to guide attention, perception, and behavior (Esghaei et al., 2022a; Fries, 2015, 2023; Jensen et al., 2007, 2012; Nobre & Van Ede, 2023; VanRullen, 2016, 2018; Weisz et al., 2014). However, the spectral signatures supporting visuospatial attention before stimulus onset remain debated (Spadone et al., 2021; Yang et al., 2025). Using MEG and a Posner-like cueing paradigm, we examined how oscillatory activity, network synchronization, and theta–gamma phase-amplitude coupling during the cue–target interval shape conscious reports of near-threshold targets.

### Temporal dissociation between beta oscillations

Results revealed a temporal dissociation in the role of beta oscillations during the cue-target interval, indicating that distinct oscillatory states support spatial attention before conscious perception (Bartolomeo et al., 2025; Liu et al., 2023; Seidel Malkinson et al., 2024).

Early in the cue–target interval, beta oscillations in the right SPL preceded successful reports of validly cued targets. This right-lateralized activity suggests spatial orienting mechanisms mediated by the SLF I operate across the visual field to prioritize relevant locations (Shomstein & Gottlieb, 2016). The absence of a similar effect in the left SPL is consistent with dominant right-hemisphere control of spatial orienting (Bartolomeo et al., 2025; Mesulam, 1981; Nobre, 2001). As a DAN node implicated in attentional shifts and rhythmic scanning (Corbetta et al., 2008; Fries, 2023), the SPL may implement beta-mediated sampling of the cued location (Behrmann et al., 2004; Buschman & Miller, 2007; Vandenberghe et al., 2012).

Beta rhythms have been linked to top-down feedback and goal-directed processing. Alpha and beta activity may reflect feedback signaling, distinguishing them from higher-frequency feedforward processes (Xiong et al., 2023). Converging causal evidence supports this view: lesions to frontoparietal attention networks produce periodic attention deficits aligned with beta rhythms (Raposo et al., 2023), and prefrontal beta coherence modulates exogenous attentional selection under conditions of stimulus conflict (Dubey et al., 2023). Consistent with primate findings of beta coherence between parietal and frontal regions during top-down attention (Buschman & Miller, 2007), SPL beta activity here index salience-based prioritization before target onset (Shomstein & Gottlieb, 2016).

Prior work emphasized alpha power as an index of local excitability and network routing (Weisz et al., 2014; Wutz et al., 2014; Zhou et al., 2021). Our results instead suggest that cue-driven beta dynamics in the SPL reflect configuration of attentional networks during the cue-target interval. In line with Zhou and colleagues (2021), these preparatory states may bias perceptual report primarily through shifts in decision criterion. The absence of strong alpha modulation may reflect the structured attentional states induced by predictive cues, which engage beta-mediated top-down coordination rather than spontaneous alpha fluctuations.

Later in the cue–target interval, beta oscillations in a right temporo-occipital node (Sani et al., 2021) preceded reports of invalidly cued targets, consistent with a preparatory “lookout” mechanism facilitating detection of unexpected stimuli (Liu et al., 2023). This beta increase may prepare reorienting processes when cue predictions are violated, extending our previous findings of delayed frontoparietal recruitment following invalid cues (Spagna et al., 2022). The pre-target beta modulation observed here in the right TO node may represent an earlier, anticipatory marker of reorienting. Importantly, beta activity did not uniformly facilitate performance: elevated beta improved perception when reorienting was required (Seen Invalid > Seen Valid, see Figure 3g) but was detrimental when expectations were correct (Unseen Valid > Unseen Invalid). Although the TO node is not classically included in reorienting models, its right-hemisphere location and connectivity suggest participation in VAN-related preparatory processes. This view is consistent with the right-lateralization of the ventral attention network (Corbetta et al., 2008; Petersen & Posner, 2012) linked by the ventral branch of the superior longitudinal fasciculus (SLF III) (Thiebaut De Schotten et al., 2011).

Further, the temporal dissociation between beta oscillations resonates with the “temporal windows” framework (Wutz et al., 2014), suggesting that distinct oscillatory states organize perceptual processing across time. Our results identify an early parietal orienting window and a later temporo-occipital reorienting window within the cue–target interval, indicating that beta-mediated processes emerge at different moments before target onset. This pattern suggests that attentional control over conscious report operates through temporally structured oscillatory regimes rather than a single sustained preparatory state.

### Interhemispheric connectivity profiles display signatures of attention supporting conscious reports

Connectivity analyses revealed complementary temporal dynamics consistent with the Communication-Through-Coherence framework (Fries, 2015), and highlight a dual temporal role of gamma-band coherence in attentional reorienting.

Early gamma coherence between the right TO node and right MFG was greater for Unseen Invalid than Unseen Valid trials, with no significant difference between Seen conditions. This pattern suggests premature engagement of ventral attention circuitry (Corbetta et al., 2008; Liu, et al., 2023; Petersen & Posner, 2012) that interferes with perceptual access. Hence, right-lateralized early fronto-temporal synchronization may impair perceptual access by triggering premature shifts, which are detrimental regardless of cue validity.

Later in the interval, low-gamma coherence between the left TPJ and right visual cortex was highest for Seen Invalid trials, while the reverse pattern was observed for Unseen Valid versus Unseen Invalid trials. This interhemispheric coupling may prepare successful reorienting when expectations are violated. This interpretation aligns with clinical and tractography evidence showing that interhemispheric connectivity plays a key role in spatial attention and recovery from neglect (Lunven et al., 2015, 2019). Specifically, pre-target left prefrontal activity may lead to neglect-related, left-sided omissions (Rastelli et al., 2013) when it is not properly controlled by right-hemisphere attention networks (Bartolomeo et al., 2025). More recently, Kaufmann et al., (2024) showed that interhemispheric connectivity plays a critical role, particularly in patients with severe initial impairments, and proposed a dynamic framework in which interhemispheric interactions support recovery depending on structural constraints. Our findings extend this framework to normal perception, suggesting that interhemispheric gamma coherence may facilitate preparatory reorienting.

Together, these results indicate a temporal shift in connectivity during the cue–target interval: early fronto-temporal coupling predicts perceptual failures, whereas later posterior interhemispheric coherence supports conscious report during invalid cueing. These dynamics align with proposals that top-down attention operates through frequency-specific coupling mechanisms (Esghaei et al., 2022b), and suggests that attention operates through temporally structured windows (Weisz et al., 2014; Wutz et al., 2014) that either facilitate or constrain conscious access depending on the match between expectations and sensory input.

### Left-lateralized prefrontal nodes support conscious reports of future targets

Across analyses, left-lateralized prefrontal regions consistently supported conscious report.

First, late gamma activity in the left MFG was greater for Seen trials, suggesting a role in conscious access. This aligns with evidence implicating the left prefrontal cortex in modulating visual awareness. For example, inhibitory transcranial magnetic stimulation over the left frontal eye field impaired conscious report, likely via attentional orienting mechanisms (Chica, et al., 2013). As noted above, low-beta activity in the left frontal cortex was found to predict left-sided omissions in neglect patients (Rastelli et al., 2013). In the healthy brain, left prefrontal activity can bias conscious report by shaping perceptual decision-making (Heekeren et al., 2004).

Theta–gamma PAC in the left IFG also increased before Seen trials, indicating cross-frequency coordination supporting perceptual access. Further, late-occurring coherence between the left IFG and right MFG in the low gamma band increased during conscious reports, suggesting that bilateral prefrontal communication underpins access to near-threshold stimuli (Martín-Signes et al., 2024). Consistent with models in which theta rhythms organize gamma bursts to regulate information processing (Jensen & Colgin, 2007), this mechanism may enhance representation of near-threshold stimuli. Consistent with this framework, evidence from intracranial recordings shows that the amplitude of fast gamma activity is systematically modulated by the phase of slower theta oscillations, indicating a form of cross-frequency coupling through which theta rhythms coordinate large-scale neuronal ensembles to shape information processing (Jensen & Colgin, 2007). These findings suggest that prefrontal PAC contributes to temporally structured attentional control not only in the post-target period (Canolty & Knight, 2010; Szczepanski et al., 2014) but also before target occurrence.

### Limitations

Our ROI-based approach may have overlooked broader whole-brain dynamics (e.g., the pulvinar; Fiebelkorn et al., 2019), and contributions from other frequency bands (Asanowicz et al., 2023; Landry et al., 2024; Meehan et al., 2021). The fixed 300-ms cue–target interval may have limited detection of slower attentional rhythms and their associated gamma modulations (Chen & Gong, 2022; Jensen et al., 2012; Senoussi et al., 2019; Yang et al., 2025). Finally, although our sample size (N = 14) is within the typical range for MEG prestimulus studies (e.g., Wutz et al., 2014, N = 14; Weisz et al., 2014, N = 12), replication in larger cohorts will help strengthen generalizability.

## Conclusion

In summary, our results reveal hemisphere-asymmetric preparatory states that shape conscious perception, refining models linking attention and awareness (Cogitate Consortium et al., 2025). In particular, our results highlight hemispheric asymmetries in attentional activity preceding conscious report—a feature rarely considered in current models of consciousness (Bartolomeo et al., 2025; Seth & Bayne, 2022). Future research should test whether these oscillatory dynamics also support internally guided attention, helping determine whether conscious access relies on a shared attentional infrastructure across external and internal representations (Liu, et al., 2023; Nobre & Van Ede, 2023; Spagna et al., 2021, 2024).

## Acknowledgements

We would like to thank Dimitri Bayle, Ph.D. and Ana B. Chica, Ph.D. for help in designing the experiment and collecting the data, Brigitte C. Kaufmann, Ph.D., Monica N. Toba, Ph.D., and two anonymous reviewers for providing insightful comments and suggestions. The work of P.B. is supported by the Agence Nationale de la Recherche through ANR-16-CE37-0005 and ANR-10-IAIHU-06, and by the Fondation pour la Recherche sur les AVC through FR-AVC-017. The work of J.L. is supported by specific funding from INSERM, Paris Brain Institute, and Dassault Systèmes. Author Contribution: literature search: all authors; project administration: P. B. Data Analysis: A. S. Writing - review & editing: All authors.

---

**Supplementary Figure 1.**
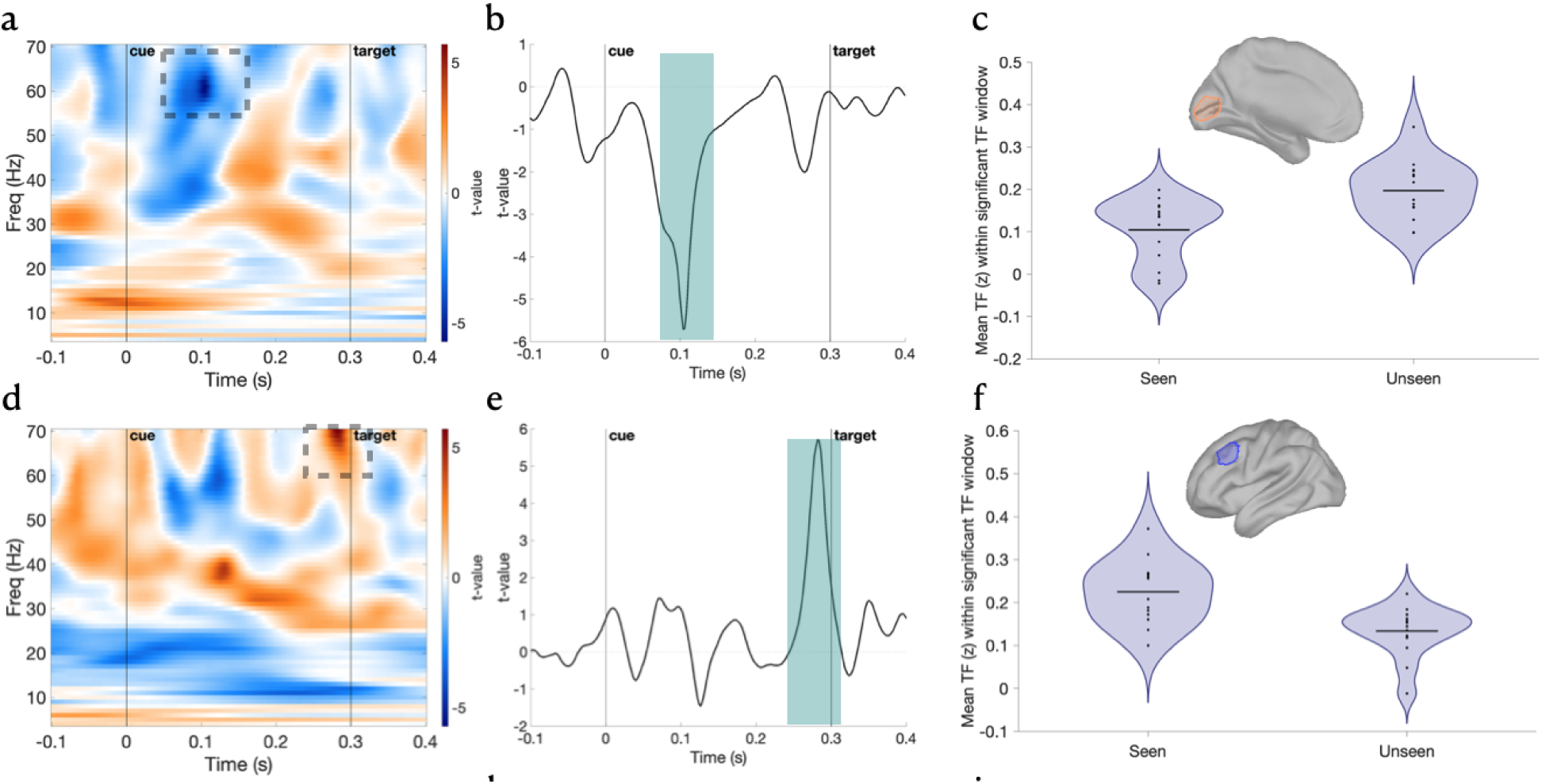
Additional gamma band effects of perceptual report (max-T permutation, 10,000 iterations, two-tailed, FWER .05; *p* < .05). **(a–c) Left medial visual cortex (Report main effect, gamma-band)**. (a) Time–frequency *t*-map shows significantly reduced gamma-band activity (∼61 Hz) at ∼104 ms post-cue for Seen vs. Unseen trials. (b) Temporal evolution of the *t*-values at the peak frequency (61 Hz); the shaded region denotes the significant time window. (c) Violin plot of individual mean amplitude values within the significant time–frequency tile: mean = 0.105 (SD = 0.074) for Seen, 0.197 (SD = 0.069) for Unseen (Cohen’s *d* = −1.53). Inset shows the cortical mask of the ROIs in the left medial visual cortex. **(d–f) Left middle frontal gyrus (Report main effect, gamma-band)**. (d) Time–frequency t-map showing significantly enhanced gamma-band activity (∼70 Hz) at ∼282 ms post-cue for Seen vs. Unseen trials. (e) Temporal evolution of the *t*-values at the peak frequency (70 Hz); shaded region highlights the significant effect window. (f) Violin plot shows group-level difference: mean = 0.225 (SD = 0.074) for Seen, 0.134 (SD = 0.059) for Unseen (Cohen’s *d* = 1.53). Inset shows the cortical mask of the ROIs in the left middle frontal gyrus. Dashed rectangles in (a) and (d) mark significant time–frequency tiles identified by the max-T test. Shaded areas in (b), (e) correspond to the significant time point.

**Supplementary Figure 2.**
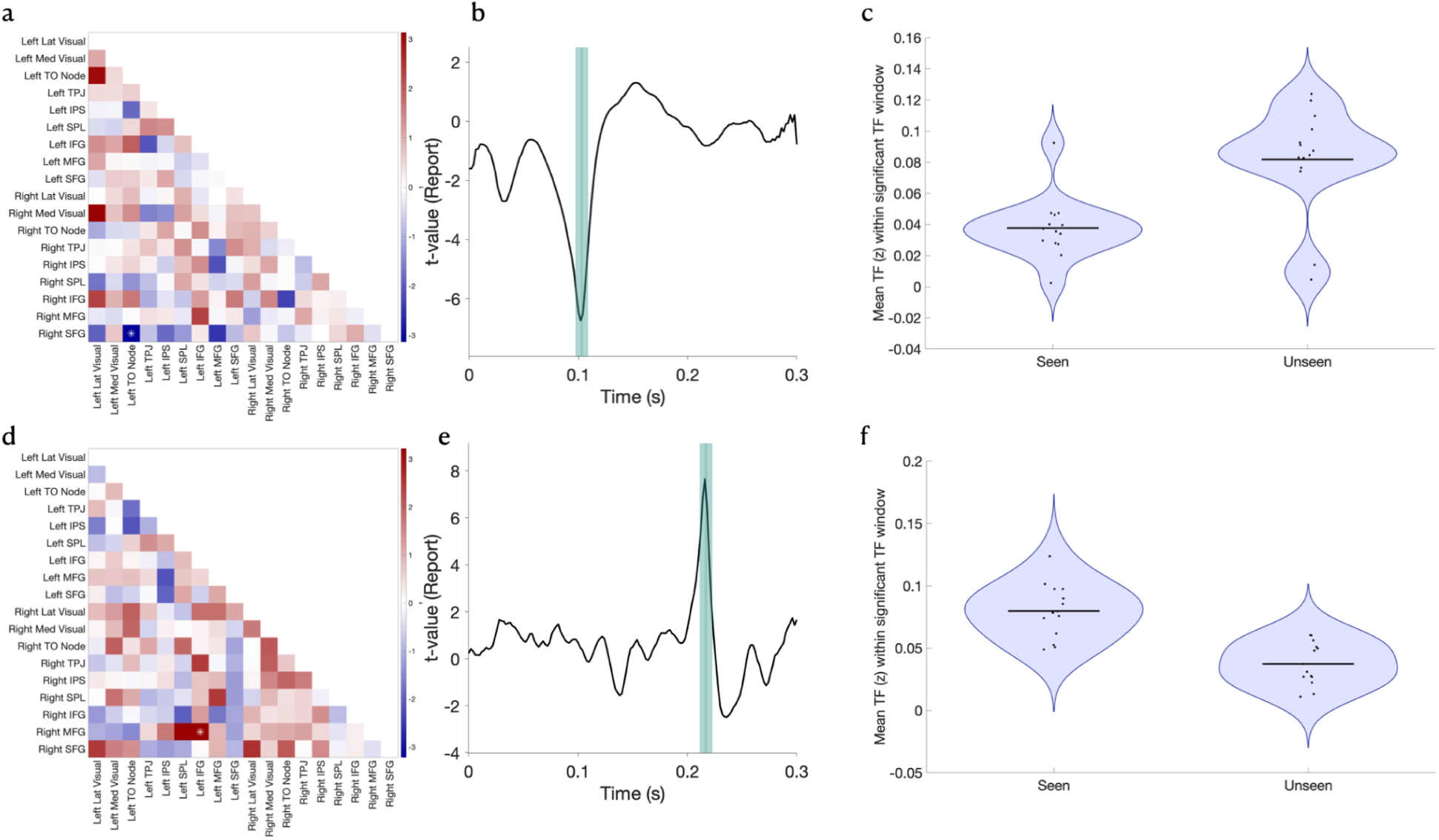
Distributed modulation of fronto-parietal coherence by visual awareness and attention. Panels show effects from time–frequency analysis of ROI-to-ROI coherence (z-transformed), summarized across the significant time–frequency windows for each contrast (max-T permutation, 10,000 iterations, two-tailed, FWER .05; *p* < .05). The full list of ROI names and order is provided in the Supplementary Table 1. (a, d) Lower-triangle connectivity matrices display t-values for each pairwise connection across 18 ROIs. Warm/cool colors indicate stronger/weaker coherence in the contrast (main effect of Report or Report x Validity interaction). (b, e) Temporal evolution of t-values at the significant connection. The shaded area indicates the time window of the significant effect. (c, f) Violin plots summarize the distribution of mean coherence values across subjects for the ROIs and time–frequency window identified in each contrast. **Top row (a–c): Right SFG – Left TO Node (beta band): main effect of Report.** A significant effect was observed in the beta band (∼102 ms post-stimulus), with coherence significantly greater for Unseen trials (mean = 0.082, SD = 0.034) than for Seen trials (mean = 0.038, SD = 0.020). This pattern indicates enhanced fronto-occipital beta-band coupling when visual stimuli fail to reach conscious awareness. **Middle row (d–f): Right MFG – Left IFG (low gamma band): main effect of Report.** A second Report-related effect emerged at ∼216 ms in the low gamma band. Coherence between the Right MFG and Left IFG was significantly stronger during Seen trials (mean = 0.080, SD = 0.022) compared to Unseen trials (mean = 0.037, SD = 0.017), reflecting enhanced prefrontal gamma-band communication during conscious visual perception.

**Supplementary Figure 3.**
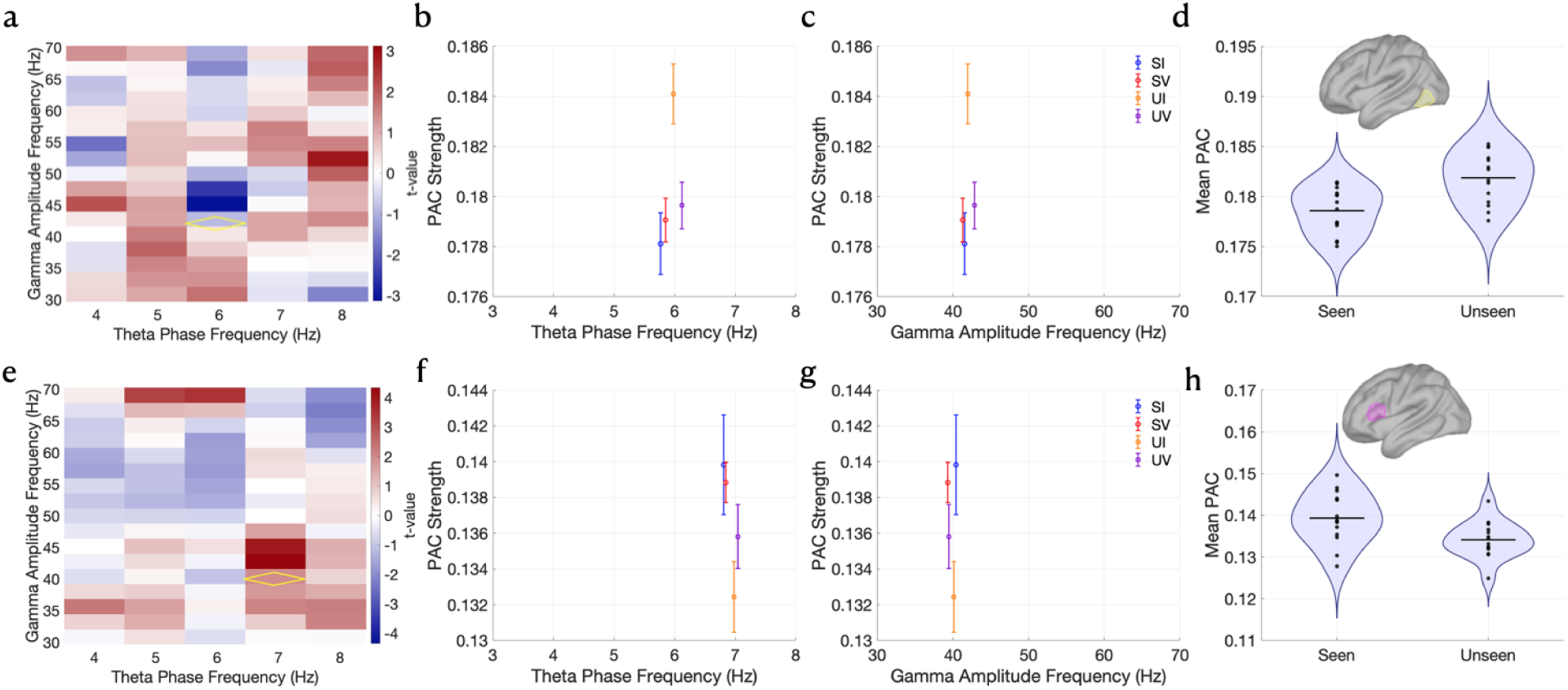
Additional PAC effects across fronto-temporo-occipital regions. Each row illustrates significant PAC modulations identified using a non-parametric max-T procedure (max-T permutation, 10,000 iterations, two-tailed, FWER .05; *p* < .001). Columns show: **(a, e)** T-value maps across theta phase (x-axis) and gamma amplitude (y-axis) frequencies, with the significant tile outlined in yellow; **(b, f)** PAC strength as a function of theta phase at the significant phase frequency; **(c, g)** PAC strength as a function of gamma amplitude at the significant amplitude frequency; **(d, h)** Violin plots showing the distribution of mean PAC values across the significant tile for each condition, with individual participants plotted as black points and horizontal bars marking condition means. **Top row (a–d): Left temporo-occipital (TO) node — main effect of Report.**A significant effect was observed at 6 Hz (theta phase) and 42 Hz (gamma amplitude). PAC strength was significantly greater for Unseen trials (mean = 0.183, SD = 0.005) compared to Seen trials (mean = 0.179, SD = 0.006). The violin plot in (d) summarizes this difference, indicating stronger theta–gamma coupling in the left TO during trials where stimuli were not consciously perceived. Inset shows the cortical mask of the left TO node ROIs. **Middle row (e–h): Left inferior frontal gyrus (IFG) — main effect of Report.** A significant effect was found at 7 Hz (theta phase) and 40 Hz (gamma amplitude). PAC strength was significantly greater for Seen trials (mean = 0.140, SD = 0.007) relative to Unseen trials (mean = 0.135, SD = 0.007). The violin plot in (h) illustrates this report-related enhancement, suggesting that frontal PAC increases during conscious access. Inset shows the cortical mask of the left IFG ROI.

**Supplementary Table 1.** Detail regarding how the 18 ROIs were created, including seed vertex and number of vertices included, surface area and Brodmann area number. LH = left hemisphere; RH = right hemisphere.

| ROIs | Seed Vertex | Seed Vertex Coords. | Region | BA | Number of Vertices | Area (cm<sup>2</sup>) |
| --- | --- | --- | --- | --- | --- | --- |
| Left Lateral Visual | 3595 | -23 -93 5 | LO | BA18 | 120 | 16.35 |
| Left Medial Visual | 4601 | -11 -83 9 | LO | BA17 | 120 | 15.91 |
| Left Temporo Occipital Node | 4018 | -42 -67 -6 | LT | BA37 | 120 | 15.24 |
| Left TPJ SLF 3 | 1926 | -56 -47 26 | LP | BA40 | 120 | 18.65 |
| Left IPS SLF 2 | 2056 | -42 -67 53 | LP | BA39 | 120 | 20.35 |
| Left SPL SLF 1 | 3238 | -23 -70 55 | LP | BA7 | 120 | 17.01 |
| Left IFG | 5676 | -57 23 20 | LF | BA44 + BA45 | 120 | 18.94 |
| Left MFG | 1558 | -38 24 40 | LF | BA46 | 120 | 15.73 |
| Left superior Frontal SLF1 | 6197 | -24 27 59 | LF | BA8 + BA9 | 120 | 17.71 |
| Right Lateral Visual | 10978 | 30 -92 7 | RO | BA18 | 120 | 15.71 |
| Right Medial Visual | 11093 | 13 -82 10 | RO | BA17 | 120 | 16.51 |
| Right Temporo Occipital Node | 12250 | 43 59 -6 | RT | BA37 | 120 | 16.13 |
| Right TPJ SLF 3 | 13831 | 55 -33 36 | RP | BA40 | 120 | 20.08 |
| Right IPS | 13907 | 53 -57 50 | RP | BA39 | 120 | 17.48 |
| Right SPL | 13496 | 28 73 54 | RP | BA7 | 120 | 17.98 |
| Right IFG SLF3 | 8325 | 57 31 11 | RF | BA44 + BA45 | 120 | 16.57 |
| Right MFG SLF 2 | 12960 | 38 30 32 | RF | BA46 | 120 | 17.49 |
| Right superior Frontal SLF1 | 9031 | 22 22 64 | RF | BA8 + BA9 | 120 | 17.07 |

Note: Coherence was computed across all pairwise combinations of the 18 anatomically defined ROIs.

